# Phylodynamics of Somatic Evolution: A Likelihood-Based Approach for Cellular Reproduction

**DOI:** 10.1101/2024.12.05.627005

**Authors:** Tobias Dieselhorst, Johannes Berg

## Abstract

Understanding the evolutionary dynamics of cell populations requires models that link observed phylogenetic patterns to the underlying processes of cell division, death, and mutation. Classical phylodynamic inference methods—–developed primarily for macroevolutionary settings—assume that mutations accrue in calendar time and often rely on a molecular clock. Here, we introduce a framework that ties mutations to discrete birth (division) events. In this setting, mutations accumulate via a compound Poisson process, capturing both visible and hidden cell divisions within the reconstructed phylogenetic tree. We present a computationally efficient dynamic programming algorithm to compute the likelihood based on tree topologies with associated mutations, integrating over latent variables such as branch durations and unobserved cell divisions. Our method is applicable to large-scale single-cell datasets, and we demonstrate its utility on simulated data and on single-cell phylogenies of haematopoietic stem cells.

## 1 Introduction

Evolutionary analysis seeks to infer the past dynamics of a population, or several populations, on the basis of extant data. A wide range of concepts and tools have been developed to determine the evolutionary events that shaped a population, ranging from the problem of inferring rates of birth and death of species to the timing of bifurcations in trees (Nee et al. 1994; Maddison et al. 2007; Stadler 2013; Sagulenko et al. 2018; Bouckaert et al. 2019; MacPherson et al. 2022).

These inference schemes are based on specific models of (i) the underlying population dynamics generating the population and hence the reconstructed phylogenetic tree and (ii) models specifying the mutations along a phylogenetic tree. The first model typically involves a birth–death process with birth and death rates that may be homogeneous or heterogeneous across the tree (Stadler 2013; MacPherson et al. 2022). For the second model, the standard choice is the molecular clock hypothesis, which assumes that mutations accumulate at some rate, constant or variable (Drummond et al. 2006), per unit time. This approach is well-suited to a macro-evolutionary setting, where birth and death describe the speciation and extinction of species over long time-scales and many generations, and mutations arise through the procreation of individuals and the fixation of these mutations in a population.

Here, we develop an approach to phylodynamic modelling and inference where mutations are tied to the divisions of individual cells, rather than the passage of time. This link between cell divisions (birth events) and mutations is specific to cellular reproduction, where bifurcations in the phylogenetic tree correspond to divisions of individual cells, rather than speciation events of entire populations. In this case, the mutation statistics can be affected by individual birth events along a branch. Some of these birth events correspond to bifurcations of the phylogenetic tree and are easily detected. Other birth events produce a lineage that eventually dies out, and these events are not directly seen in the phylogenetic tree, see Figure 1. Yet these ‘hidden’ birth events contribute to the accumulation of mutations. Combining the number of hidden birth events along a branch with the number of mutations at each birth event leads to a compound statistics for the number of mutations along each branch. Specifically, we find that the standard Poisson statistics arising in constant-rate molecular clock models is replaced with the so-called compound Poisson statistics, leading to an increased variance arising from discrete reproductive events along branches compared to a molecular clock. Previous macroevolutionary models introduced compound Poisson statistics in an ad hoc manner to relax the molecular clock assumption (Takahata 1987; Huelsenbeck et al. 2000). In contrast, we show that this compound structure emerges naturally from the dynamics of cellular reproduction.

**Figure 1:**
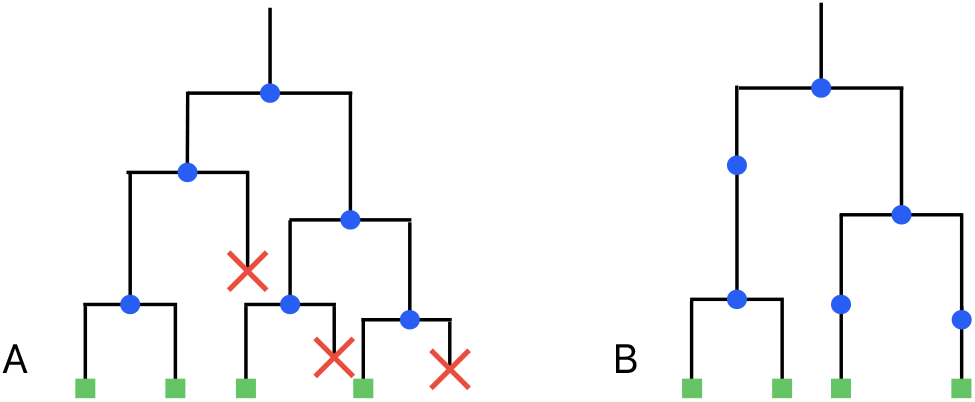
Comparison of a birth–death tree and the corresponding reconstructed phylogenetic tree. (A) A birth–death process of individuals with time running downwards consists of birth events (blue circles) and death events (red crosses) and produces some individuals (green squares) that are alive at the time of observation. Each birth event produces a bifurcation in the birth–death tree. (B) The corresponding phylogenetic tree reconstructed from extant individuals only. Birth events are again indicated by blue circles and can now occur *within* phylogenetic branches (‘hidden’ birth events), as well as at all bifurcations of the phylogenetic tree.

Our inference method is designed to handle large-scale single-cell datasets, which are derived from populations of cells undergoing division and death. Examples include the spread of bacteria, tissue development, or the uncontrolled growth of tumor cells. In order to keep the computations simple, we restrict the discussion to the infinite sites model. Under this model, the topology of the phylogenetic tree and the ancestral states are straightforward to infer. To facilitate the efficient computation of the likelihood of a tree (specified by its topology and the mutations along each of its branches), we use a dynamic programming approach. This involves iteratively integrating over all branch lengths (in calendar time) and summing over the number of cell divisions per branch. The algorithm achieves running times of just seconds for hundreds of samples and scales linearly with size of the tree.

In Section 2.1, we recapitulate the dynamics of a birth–death process and establish the statistics of mutations along tree branches. We introduce our algorithm to compute the likelihood of the corresponding model parameters and test its capabilities on simulated data. In Section 2.2, we generalize these results to trees reconstructed from a finite fraction of sampled individuals. We apply our method to single-cell data of haematopoietic stem cells (Mitchell et al. 2022) in Section 2.3, analyzing prenatal haematopoiesis as well as detecting distinct dynamics in different clades found in an adult donor.

## 2 Results

### 2.1 Parameter inference from mutations along branches

We consider the growth of a population under a constant-rate birth–death process. Without loss of generality, we chose the unit of time such that the birth rate is one and denote the rate of cell death (relative to the birth rate) by *q <* 1. By convention, time runs backwards into the evolutionary past from the time of observation. From the reconstructed phylogenetic (topological) tree and the mutations along its branches, we aim to infer the underlying phylodynamic model. Such a model consists of two components: The birth–death process (described by the rates of cell division and cell death), and the statistics of mutations arising at each cell division.

The first component, the birth–death process, is characterized by the joint statistics of the length *τ* of a branch in calendar time and the number of generations *i* along it. The following distribution combines these two measures for a clade starting with a single individual at time *τ*_*s*_ in the past: the joint probability that the individual has undergone *i* birth events by time *τ*_*e*_ = *τ*_*s*_ − *τ* and is alive at time *τ*_*e*_, but all *i* lineages generated between *τ*_*s*_ and *τ*_*e*_ have died out by the time of sampling is given by

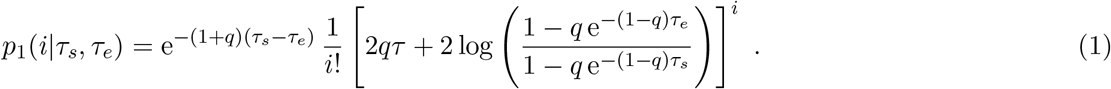

The fate of the particular individual alive at time *τ*_*e*_ after that time is not specified. A derivation can be found in Supplementary Information B. The number of generations *i* along a branch starting at *τ*_*s*_ and ending at *τ*_*e*_ is Poisson distributed with a mean that depends non-linearly on the age of the branch (Bokma et al. 2012).

The second component concerns the statistics of the mutations occurring at each birth event (cell division). Here, we consider the number of mutations per cell division to be Poisson distributed with mean *µ*, so the number of mutations along a branch with *i* generations along it plus one additional generation founding the branch takes the value *m* with probability

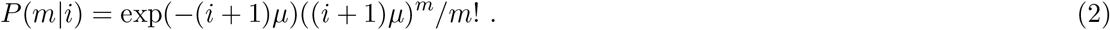

Combining this Poisson distribution with the (also Poissonian) distribution of the number of generations (1) along a given branch, the number of mutations along the branch follows a compound Poisson distribution. Compound Poisson statistics are generated by a sum of independent, identically distributed random variables, in our case the number of mutations per cell division, where the number of random variables is itself a Poisson distributed random variable (in our case the number of generations *i*). The compound Poisson distribution has a larger variance than a simple Poisson distribution; Figure 2 shows two examples. For a sufficiently large number of mutations per generation, one can see the effects of individual cell divisions along a branch in the form of oscillations. For details on the compound Poisson distribution, see Supplementary Information C.

**Figure 2:**
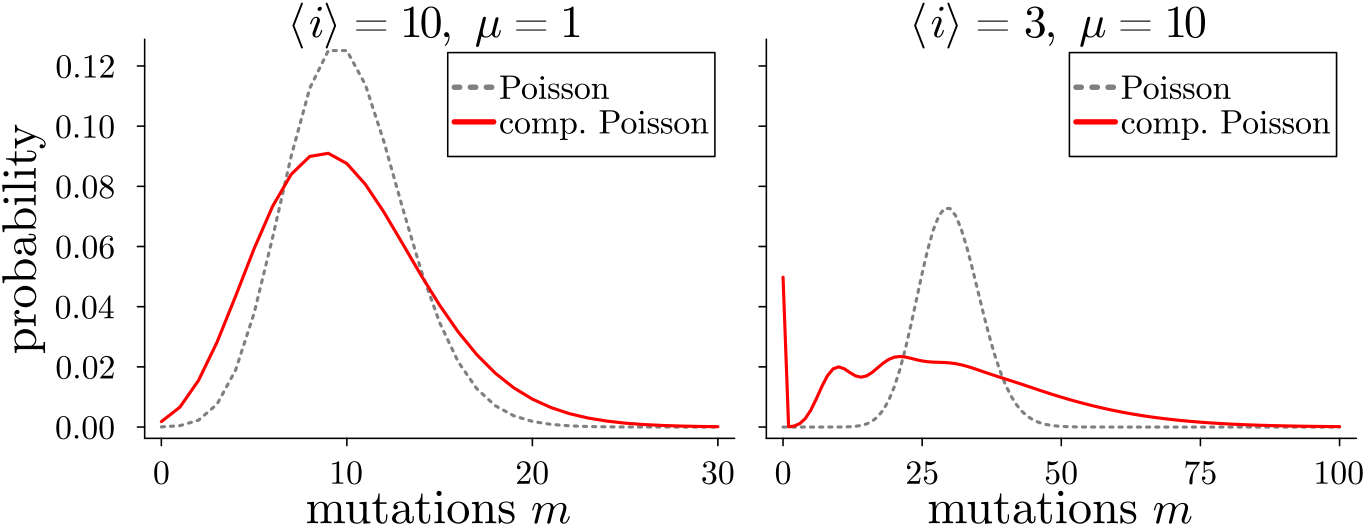
Compound Poisson distributed number of mutations. The distribution of the numbers of mutations along a branch follows a compound Poisson statistics arising from the discrete number of generations along a branch. The red solid lines show the distributions of the number of mutations along branches with mean number of generations ⟨*i*⟩ and mean number of mutations per generation *µ*. They are compared to Poisson distributions (grey dashed lines) with the same mean ⟨*i*⟩ *µ* as they would arise from a molecular clock. Continuous lines are drawn for visibility. See Supplementary Information C for details.

We now combine these two components of a phylodynamic model to construct a likelihood function based on the probability that a tree with the empirically observed numbers of mutations along its branches arises under a birth–death process and the mutation statistics (2). Branch lengths in calendar time and the number of birth events along each branch on the other hand are unobserved variables. In contrast to macroevolutionary data, in single-cell data there are often only few mutations along a branch, so branch lengths cannot be determined precisely. They act as latent variables in this inference problem. To sum over these latent variables, we use a simple iterative procedure, dynamic programming, in the spirit of Felsenstein’s recursive algorithm (Felsenstein 1981): we iteratively sum over both the times of bifurcations and the number of generations along each branch. The same iterative approach applied to bifurcation times is used in the *TreeTime* algorithm by Sagulenko, Puller, and Neher (Sagulenko et al. 2018).

*P* (*ν*|*τ*_*s*_) denotes the probability that the part of the phylogenetic tree below a given branch *ν* (starting at the top of the branch which emerged at time *τ*_*s*_) arose under a particular phylodynamic model. In this definition, the phylogenetic tree is specified by its topology and the number of mutations along the branches. The phylodynamic model is specified by the relative death rate *q* and the mean number *µ* of mutations per cell division. For a pendant branch (a branch starting at *τ*_*s*_ and ending in a leaf) with *m* mutations along it, *P* (*ν*|*τ*_*s*_) follows from the probability that a clade starting at time *τ*_*s*_ runs until time zero with *i* generations along it, but has no further extant offsprings (equation (1) with *τ*_*e*_ = 0). From the distribution of the number of mutations along a branch with *i* generations (2) we directly obtain

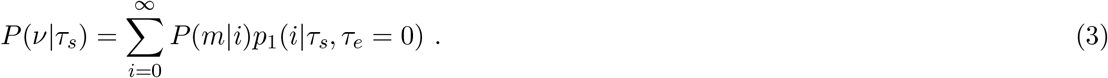

On the other hand, for an internal branch *ν* with *m* mutations along it and the two branches *ν*^*′*^ and *ν*^*′′*^ emanating from it we have

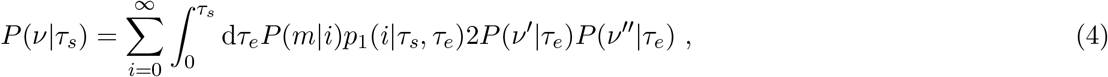

where we integrate over the time *τ*_*e*_ of the end of branch *ν* (and therefore the beginning of branches *ν*^*′*^ and *ν*^*′′*^). The factor of two arises as either of the two clades arising at *τ*_*e*_ can be *ν*^*′*^ or *ν*^*′′*^.

The likelihood for the model parameters given mutations along the tree can thus be evaluated iteratively, integrating over the calendar times of internal nodes and the number of generations along each branch in a computationally efficient manner. We start with (3) evaluated for all pendant branches, and then evaluate (4) for all branches *ν* whose *P* (*ν*^*′*^|*τ*) and *P* (*ν*^*′′*^|*τ*) have already been evaluated for their descendant branches *ν*^*′*^ and *ν*^*′′*^. From the last two remaining branches *ν*^*′*^ and *ν*^*′′*^ emanating from the most recent common ancestor, we finally obtain the likelihood of the model parameters given the observed tree (topology and number of mutations along branches)

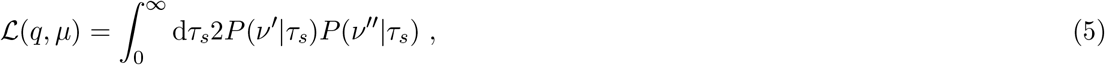

where a uniform prior has been used for the time of origin of the tree (the time at which *ν*^*′*^ and *ν*^*′′*^ arise). This improper prior could be replaced by a proper prior by using an upper limit on the time of origin. Under the conditions considered here (fixed rate of birth and a rate of death that is smaller than the birth rate), taking this upper limit to infinity yields the result from the improper prior.

To test this inference scheme, we generate a phylogenetic tree from a population grown from a single individual to size *N* = 300 (stopping when reaching this size) at *q* = 3*/*4 and *µ* = 1. Figure 3A shows the likelihood landscape defined by (5). The integrals in (4) and (5) were performed numerically using the trapezoidal rule on a grid dividing the interval [0, 2 log(*N*)*/*(1 −*q*)] into 1000 subintervals, thus setting the upper limit of the age of the tree to twice its expected value. The upper limit of the sums over generations *i* was set to 50. The computation of a single likelihood took about 2.5 to 3 seconds for a tree of *N* = 300 leaves (4.4GHz clock speed) and can be sped up by using fewer subintervals for the numerical integration. The running time scales as the second power of the number of subintervals and linearly with the upper limit on the number of generations.

**Figure 3:**
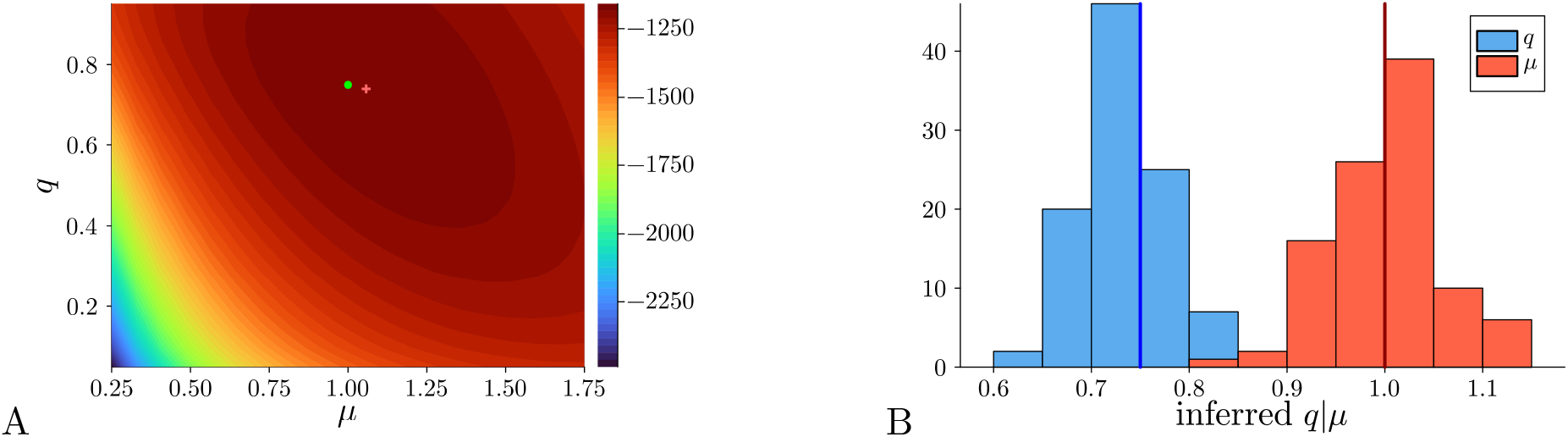
Inference using the likelihood of the birth–death process. (A) A population was grown from a single individual to *N* = 300 individuals with *q* = 3*/*4 and *µ* = 1 and the likelihood (5) for the resulting tree (defined by topology and mutations along branches) was computed iteratively as described in the text. The maximum of the log-likelihood near *q* = 0.74 and *µ* = 1.06 is marked with a red cross, the underlying parameters are marked with a green circle. (B) We grew 100 populations in the same way as in (A) and show the resulting histogram of the inferred values of *q* (left, in blue), and *µ* (right, in red) based on the likelihood (5). The underlying values *q* = 3*/*4 and *µ* = 1 are marked with blue and red vertical lines, respectively.

Figure 3B shows the histogram of parameters inferred from 100 different populations grown in the same way. As expected, the inferred values fluctuate from run to run around mean values close to the underlying values. Supplementary Information F shows that a wide range of parameters can be recovered in this way and explores how the inference improves with increasing tree size. We note that to exactly simulate the statistics conditioned on a given population size, one would have to include in the simulation also instances where this size is reached for the second, third, … time. This is subtly different from the choice here, where the population reaches the given size for the first time, see (Hartmann et al. 2010) for a discussion. For supercritical growth and large population sizes, we expect any differences between the two statistical ensembles to be minor, as fluctuations around the mean size are small.

### 2.2 Finite sampling

In most applications, a phylogenetic tree is not based on the entire extant population at the time of observation but on a subset of sampled individuals or cells. Reconstructing the phylogenetic tree from a subset of individuals changes both the statistics of branch lengths (Stadler 2010) and the number of generations along branches. Hence, sampling must be accounted for in the likelihood function.

In this section, we follow a standard model of sampling (Yang and Rannala 1997; Stadler 2010; Stadler and Steel 2012), where extant individuals are selected independently at the time of observation with equal probability *ρ*. The key property of a phylogenetic branch then becomes that all extra offspring arising in birth events along that branch either have their clade die out by the time of observation (as before), or, alternatively, no extant element of that clade is sampled.

The probabilities that a birth–death process starting with a single individual at time zero has no *sampled* offspring or a single sampled offspring, respectively, have been calculated by Yang and Rannala (Yang and Rannala 1997) as

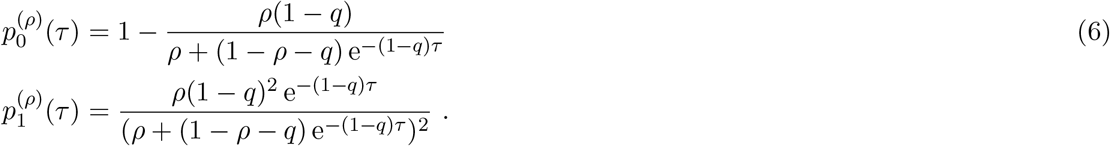

These results (6) for the probability that a clade has no sampled offspring can be used to generalize the distribution *p*_1_(*i*|*τ*_*s*_, *τ*_*e*_) to the case of finite sampling, following the steps in Appendix B. The result is

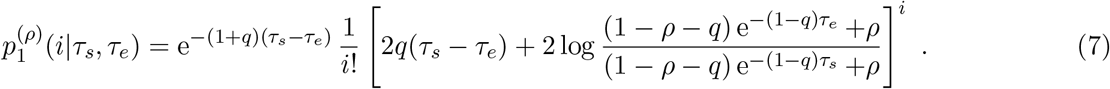

Using this equation for the likelihood computation in Section 2.1 incorporates the effect of finite sampling into the inference scheme. In the equation for pendant branches (3) an additional factor of *ρ* describes the probability that the extant individual is sampled.

In applications of single-cell sequencing such as somatic evolution or evolution of a pathogen within a host, the population size and hence the sampling probability *ρ* is generally easier to estimate than for populations of individuals in the wild. However, in principle, *ρ* can also be inferred along with the other parameters *q* and *µ*, see Supplementary Information E.

A point that is not related to sampling but has a similar effect on the tree is sequencing noise. In particular in single-cell sequencing, some of the mutations found in the samples will be artifacts produced by sequencing errors. These also lengthen the pendant branches. By incorporating sequencing noise into the likelihood (5), the level of noise can be inferred from the tree. Details are in Supplementary Information G.

### 2.3 Application to single-cell data of haematopoietic stem cells

#### 2.3.1 Prenatal haematopoiesis

As a first application, we use previously published whole-genome sequencing data of single haematopoietic stem cells (HSC) taken from the cord blood of neonates (Mitchell et al. 2022). A tree based on sequencing data from 216 HSC from donor CB001 is depicted in Figure 4A with branch lengths corresponding to the number of mutations. We estimate the sampling probability to be *ρ* = 0.01, based on an order-of-magnitude estimate of 2 *×* 10^4^ to 2 *×* 10^5^ HSC in the donor blood of adults (Mitchell et al. 2022). The maximum number of mutations along a branch is 62 found on a pendant branch. In general, pendant branches carry much larger numbers of mutations than the tree’s interior, where most branches only carry a few mutations. This agrees qualitatively with the results of (Dieselhorst and Berg 2024) for the relative lengths of pendant versus interior branches under finite sampling.

**Figure 4:**
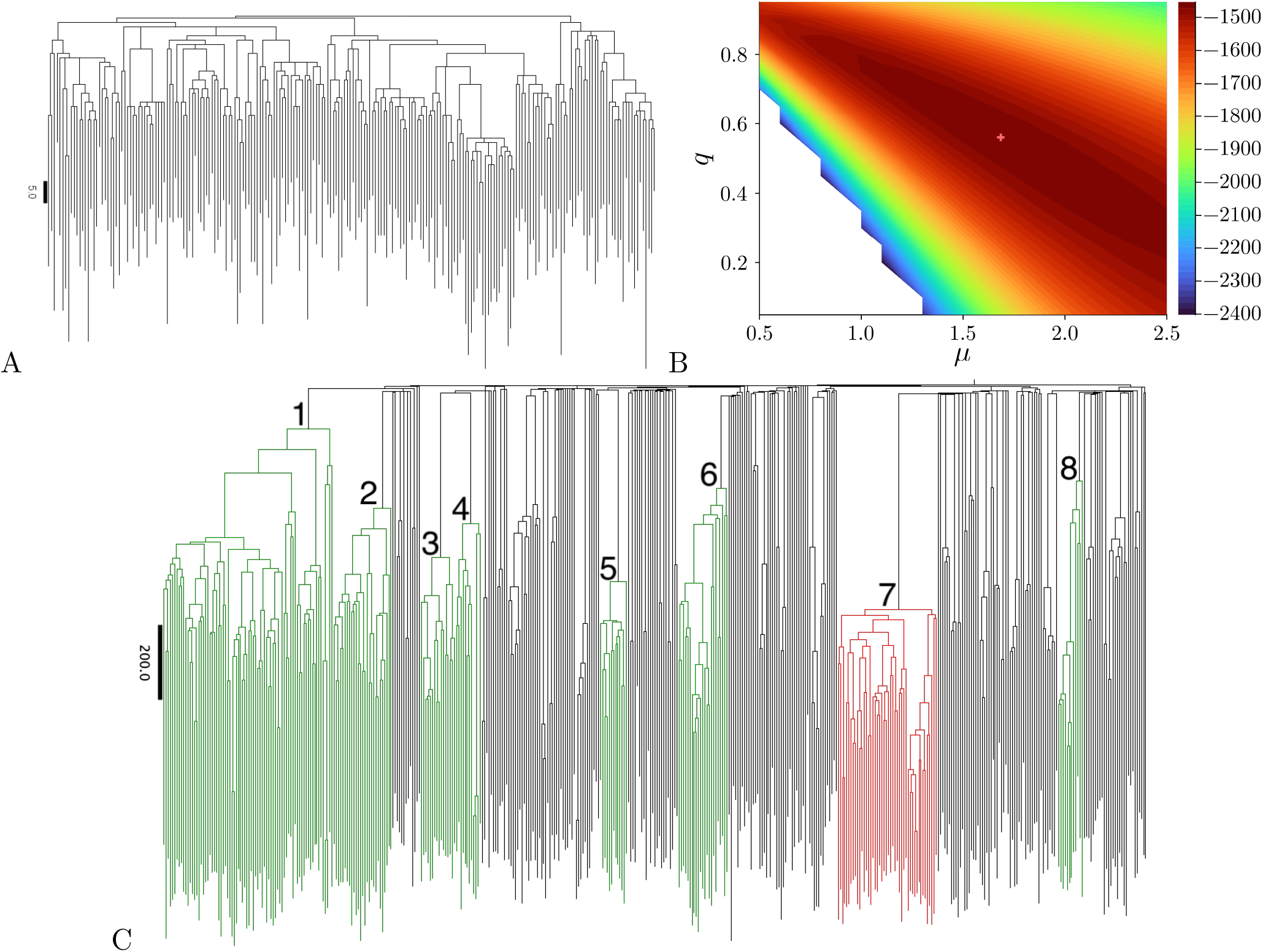
Likelihood of different trees from data. (A) The reconstructed tree from single-cell data of HSC in donor CB001 from (Mitchell et al. 2022) with branch lengths proportional to the number of mutations along them. The phylogeny is taken from (Mitchell et al. 2022). The length of the black bar on the left corresponds to five mutations. (B) The resulting log-likelihood landscape of this tree when setting the sampling probability *ρ* = 0.01. Values of the log-likelihood below − 2400 are left blank. The maximum (marked by a red cross) is found at around *q* = 0.52 and *µ* = 1.24. (C) The phylogenetic tree of donor KX004. We find strong evidence that the clade shown in red has evolved under different evolutionary parameters than the other (green) clades, see text.

HSC differentiate into different blood cells. We assume that this differentiation occurs stochastically at some rate per unit time. The inferred relative death rate will therefore correspond to the sum of the rates of cell-death *and* differentiation relative to the rate with which cells replicate. During development, the HSC population expands rapidly, so we expect that the majority of mutations arises during cell replication (Spisak et al. 2023) (as assumed in (2); to relax this assumption, see Supplementary Information G).

We compute the likelihood (5) by integrating over the time interval [0, 2 log(*N/ρ*)*/*(1−*q*)] discretized in 1000 steps and setting an upper limit of 200 for the sum over *i*. The upper limit is again twice the expected time to grow a tree with *N* samples at the given sampling probability. The results are shown for different values of the parameters in Figure 4B. The maximum of the likelihood is found to be near *q* = 0.56 and *µ* = 1.68. Given that on average interior branches carry about three mutations each this corresponds to about two birth events per branch, one of them hidden.

We also inferred the parameters for the dataset of a second neonate (donor CB002 in (Mitchell et al. 2022)), which contained 390 HSC as well as haematopoietic progenitor cells (HPC). Under the assumption that HPC behave like HSC and that the sampling probability was again *ρ* = 0.01, the maximum lies at approximately *q* = 0.64 and *µ* = 1.45.

Estimating the sampling probability *ρ* precisely is difficult. In particular for donor CB002, one can also argue for a lower value of *ρ*, as the phylogenetic tree contains both HSC and HPC and the number of HPC in humans is approximately ten times higher than that of HSC (Miao et al. 2022). We find that our parameter estimates only depend weakly on the sampling probability *ρ*. For instance, setting *ρ* = 0.001 leads to *q* = 0.52 and *µ* = 1.24 for donor CB001 and of *q* = 0.61 and *µ* = 1.03 for donor CB002, thus yielding similar relative death rates but values of *µ* which are roughly 25% lower than in the case of *ρ* = 0.01.

For comparison, we also inferred the model parameters under the assumption of a constant-rate molecular clock. We find that the log-likelihoods maxima under the two models do not differ much, meaning that for the data analyzed here either model fits the data nearly equally well. Generally the model (5) leads to somewhat higher likelihood maxima (except for case CB002 at *ρ* = 0.01). For both datasets and the range of sampling probabilities considered here, the model with a constant-rate molecular clock leads to a maximum of the likelihood at *q* = 0. The corresponding likelihood landscapes and estimates of the confidence regions around these point estimates are shown in Supplementary Information I. This result is unexpected and biologically unrealistic since differentiation at a finite rate leads to a non-zero value of *q*. The number of parameters in the two types of models — mutations at cell divisions and at a constant rate — are the same, so it appears that the model with mutations at cell divisions can explain the data better in a biologically plausible parameter regime. Changing the underlying birth–death process or considering different mutational processes may change these conclusions, see Section 3 Discussion and Supplementary Information G, respectively.

All these results were derived under the assumption of a constant-rate birth–death process (with death events corresponding to either cell death or differentiation). In the following section, we infer clade-specific evolutionary parameters in order to identify signals of selection.

#### 2.3.2 Heterogeneity between different clades in adult haematopoiesis

During adulthood, HSC and HPC continue to accumulate mutations. Some of these mutations can increase the rate of net growth. The corresponding clades can acquire further mutations that eventually, after may further steps can lead to the development of cancer. In this section we ask if in trees from adult HSC and HPC, there is evidence for cades growing at different rates from the rest, that is, heterogeneity of the model parameters across different clades.

Based on 451 HSC and HPC sampled from the bone marrow of 77 year old donor KX004, a phylogenetic tree was constructed in (Mitchell et al. 2022). We ask if clades in this phylogenetic tree differ in their parameters *q* and *µ*. As we expect a drastic difference between haematopoietic cell dynamics between an embryo and an adult, we will restrict our analysis to clades which have arisen after birth. Based on the neonate trees analyzed in the previous section (which have a maximum of 83 (CB001) and 80 (CB002) mutations between root and tip), we assume that nodes with a distance of more than 100 mutations from the root have arisen after birth of the donor. We further restrict our analysis to clades with at least 23 leaves (i.e. 45 nodes in total). The tree of donor KX004 contains eight clades fulfilling these two criteria, see Figure 4C. We analyze each of these clades independently (thus removing any correlations between the times at which the eight clades arise), as explained above. In Supplementary Information H we discuss how to identify, in a tree growing otherwise at uniform rates, a clade under selection by allowing the evolutionary parameters to change at a point within the tree.

To compute the likelihood of a single set of global parameters (*q, µ*) (i.e. the same parameters in all eight clades), we multiply the probabilities (5) for each clade, resulting in the joint probability of the observed clades. Assuming a sampling probability of *ρ* = 0.01 and using the same discretizations and intervals as in the previous Section 2.3.1, we find the likelihood maximum around *q* = 0.91 and *µ* = 10.81.

The inference from multiple clades allows for an easy detection of differences between the evolutionary dynamics in the different clades. In principle, each clade can be assigned a separate set of variables. As the optimization of such a high-dimensional problem is not feasible, we consider distinct parameters *q*_dist_ and *µ*_dist_ in just one of the eight clades. All other clades follow the model with parameters *q*_main_ and *µ*_main_. We assume a homogeneous sampling probability *ρ* throughout.

We have computed maximum likelihood estimates for eight different heterogeneous scenarios, each of them assigning distinct parameters to a different clade (and different parameters to the rest). The results are shown in Table 1. We show the likelihood ratio

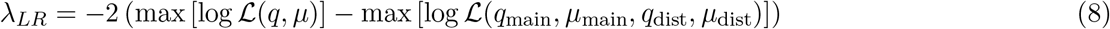

defined as the logarithm of the ratio of the maxima of the homogeneous and heterogeneous model, multiplied by a factor of two. In the limit of infinite data, *λ*_*LR*_ follows a *χ*-squared distribution (in the case here with two degrees of freedom, as this is the number of additional parameters in the heterogeneous model)(Wilks 1938). Thus, the *χ*-squared distribution provides an estimate of the *p*-value of each heterogeneous model, i.e. the probability of falsely rejecting the homogeneous model in favour of the heterogeneous model. We set a significance threshold of *α* = 0.05 to reject the homogeneous model in favour of a heterogeneous model. As we are testing eight different heterogeneous models, only the ones which have a *p*-value below the Bonferroni corrected significance value *α/*8 = 0.00625 are accepted. Only the scenario with distinct parameters in clade 7 fulfils this criterion.

Changing the sampling density from *ρ* = 0.01 to *ρ* = 0.001 does not change the clade: again only clade 7 is detected as having distinct parameters from the rest of the clades. The likelihood maximum for this clade shifts from a decrease in the relative death rate *q* to a decrease in the mutations per birth event *µ* = 5.44. However, the likelihood difference to the alternative scenario where *q* is decreased and *µ* is unchanged is insignificant, so these two scenarios cannot be resolved with the current amount of data.

Repeating the analysis with *ρ* = 0.001 and assuming same dynamics across the entire tree, the maximum likelihood estimate yields a similar relative death rate *q* = 0.90 but a lower mean number of mutations per birth event *µ* = 7.22. Note the similarity to the dependency on *ρ* in the previous analysis in Section 2.3.1. The detection of distinct dynamics does not appear to be very sensitive to the choice of *ρ*, as we again find different dynamics in clade seven with the same p-value as before. However, the inferred difference in dynamics changes, as in the case of *ρ* = 0.001 we find similar relative death rates across the tree (*q*_main_ = 0.89 and *q*_sub_ = 0.90) but fewer mutations per birth event in clade 7 (*µ*_main_ = 7.94 and *µ*_sub_ = 5.44).

**Table 1:** Maximum likelihood estimates of the parameters in eight of the clades in the phylogenetic tree from donor KX004. In each column, we consider a parameter change in one of the clades.

| **distinct clade:** | **1** | **2** | **3** | **4** | **5** | **6** | **7** | **8** |
| --- | --- | --- | --- | --- | --- | --- | --- | --- |
| $\lambda_{LR}$ | 6.17 | 1.20 | 2.27 | 1.45 | 1.74 | 0.84 | 16.23 | 2.17 |
| p-value | 0.046 | 0.549 | 0.321 | 0.484 | 0.419 | 0.657 | 0.0003 | 0.338 |
| $q_{\text{main}}$ | 0.90 | 0.91 | 0.91 | 0.91 | 0.90 | 0.91 | 0.91 | 0.91 |
| $\mu_{\text{main}}$ | 10.76 | 11.1 | 10.95 | 10.73 | 10.98 | 10.70 | 10.76 | 10.60 |
| $q_{\text{sub}}$ | 0.91 | 0.92 | 0.90 | 0.87 | 0.89 | 0.91 | 0.80 | 0.85 |
| $\mu_{\text{sub}}$ | 11.18 | 10.02 | 9.42 | 14.39 | 10.16 | 11.60 | 14.45 | 15.88 |

## 3 Discussion

Under cellular reproduction, mutations can arise at individual birth events rather than accumulating continuously over time. We have built a population dynamic framework for the phylodynamic analysis under cellular reproduction. Our approach is geared towards large-scale single-cell data: we use dynamic programming to iteratively sum over the unknown bifurcation times and the number of generations along a branch. This leads to a likelihood function that can be computed in seconds for several hundred samples and scales linearly with the number of samples.

Cellular reproduction generates phylogenetic trees which differ in two respects from their counterparts created in macroevolution. First, when mutations arise at discrete reproductive events, the number of mutations accumulated along a branch follows a compound Poisson distribution. This leads to a larger variance compared to the Poisson distribution observed when mutations arise continuously over time. Second, the mean number of mutations along a branch is not proportional to the branch length in calendar time, even when the parameters of the birth–death process are constant over time. Instead, the mean number of mutations depends non-linearly on the age of a particular branch and the birth and death rates of cells. This is because additional lineages established by birth events along a branch must die out by the time the population is sampled; otherwise, the branch terminates and two new branches begin. This condition is harder to fulfil in recent branches, reducing the number of generations per time in recent branches (Bokma et al. 2012). As a result, generations accumulate in branches at different rates in different parts of the *reconstructed* tree, even when all underlying parameters are constant through time. This can lead to errors in phylodynamic analysis whenever neutral evolution under cellular reproduction is modelled by a constant-rate molecular clock (see Supplementary Information D).

Throughout, we use the number of mutations along the branches of a tree as a basis for the likelihood. This approach can easily be extended to sequence similarity scores in order to quantify the rates of different types of mutations. We have also used the infinite sites assumption (no back mutations), so the tree topology and the characters of internal nodes (ancestral sequences) can be determined unambiguously. If this assumption breaks down, for instance in the presence of back mutations, one needs to sum both over the sequences at internal nodes (Felsenstein 1981) and potentially over tree topologies as well. This leads to a computationally harder problem, which can be addressed by Markov chain sampling (Bouckaert et al. 2019; Douglas et al. 2025). Sums over ancestral states and branch lengths can be performed by heuristics like the *TreeTime* algorithm (Sagulenko et al. 2018), which is based on alternating between the two sums. However, despite the additional computational difficulty, the statistical framework leading to the likelihood (5) generalizes directly regimes outside the infinite sites assumption. Similarly sequencing noise, in particular allelic dropout, can lead to ambiguities in the assignment of mutations to branches, which can be dealt with by summing over the different possible assignments.

Our approach differs from coalescent-based phylodynamic inference (Johnson et al. 2023), which tracks bifurcations in the tree backwards in time. The coalescent statistics depends only on the net growth rate of the population (difference between birth and death rates). However, the hidden birth events are closely related to cell death, so the coalescent cannot capture the statistics of hidden birth events. Our approach also differs from the recent analysis of punctuated evolution (Manceau et al. 2020; Douglas et al. 2025), which uses Monte Carlo Markov sampling, whereas we iteratively sum over the generations along branches/integrate over branch lengths. Punctuated evolution is a macroevolutionary model characterized by bursts of mutations at speciation events in addition to a regular molecular clock. Formally, mutations occurring at birth events can be viewed as an extreme case of punctuated evolution. Our approach also differs from inference based on the distribution of pairwise distances between the tree leaves (Werner et al. 2020) in that we compute the likelihood of the tree and use the statistics of hidden birth events arising from the birth–death process, see also (Angaji et al. 2024).

In this paper, we have focussed on the statistics of mutations that arise at cell divisions. Even in somatic evolution, mutations can arise per unit time: DNA lesions which form spontaneously might not be repaired, and thus accumulate over time. This process has been analyzed in bulk data on the basis of its mutational signature (Satake et al. 2023; Spisak et al. 2023). Under a low rate of cell division, with few replications in a given time interval, more mutations arise due to lesions which accumulate over time, whereas in fast-dividing populations, mutations arise predominantly as part of the reproductive process (Spisak et al. 2023). In Supplementary Information G we have extended our inference scheme to mixtures of mutations arising at cell divisions and mutations arising per unit time. Inferring the relative weight of these two mechanisms from data is an exiting prospect and will become feasible when larger whole-genome single-cell phylogenies become available. Furthermore, cells divide only after going through a cell cycle of finite duration. Introducing an age-dependency for birth events (Mulberry et al. 2025) is an interesting future extension of the constant-rate birth-death process currently underlying our mutation model.

## Code availability

Code is available under MIT license at https://github.com/t-dslhrst/SingleCellPhylodynamicInference.

## Acknowledgments

This work was funded by the Deutsche Forschungsgemeinschaft (DFG, German Research Foundation) grant SFB1310/2 - 325931972. Many thanks to Tibor Antal, Samuel Johnston, Joachim Krug, Martin Peifer, and Shamil Sunya’ev for discussions, and to Tanja Stadler for comments throughout and especially for helpful advice on Section 2.2. We thank Jamie Blundell, Nico Borgsmüller, Donate Weghorn, and Xu Xun for advice on single-cell data and Laura Tomás for advice on sequencing noise in Section G. We thank the computing centre of the University of Cologne RRZK for computing time on the DFG-funded clusters CHEOPS (grant number INST 216/512/1FUGG) and RAMSES (grant number INST 216/512-1 FUGG).

