## Supplementary Information for "Phylodynamics of Somatic Evolution: A Likelihood-Based Approach for Cellular Reproduction"

#### A Stochastic dynamics of a birth–death process

We model the dynamics of a population as a birth–death process with birth rate $\beta$ and death rate $\delta$ per individual and per unit time. A particular clade (one of many) begins to grow from a single individual born at time $t_0$ until time $T$, when the population is observed. We limit ourselves to the case of a growing population with $\beta > \delta$.

From a population sampled at one given time, no quantity with units of time can be inferred. A goal will be to determine the relative rate of death $q = \delta/\beta$, and without loss of generality, we set the unit of time such that the birth rate is $\beta = 1$.

The standard approach to such a birth–death process is a master equation for the size of a clade $n$ at time $t$ described by the probability $p_n(t)$ evolving under

$$\partial_t p_n(t) = \beta(n-1)p_{n-1}(t) - \beta n p_n(t) + \delta(n+1)p_{n+1}(t) - \delta n p_n(t) \ . \tag{1}$$

As initial conditions, we will assume the clade to start with a single individual at time $t = 0$, i.e. $p_1(t=0) = 1$ and $p_n(t=0) = 0$ for all $n > 1$. This equation can easily be solved using a generating function. A classic result by Kendall (Kendall 1948) gives the probability that the clade has died out by time $T$

$$p_0(T) = q\frac{1-\mathrm{e}^{-(1-q)T}}{1-q\,\mathrm{e}^{-(1-q)T}} \tag{2}$$

and the probability that there is exactly one surviving offspring by time $T$

$$p_1(T) = \frac{(1-q)^2\,\mathrm{e}^{-(1-q)T}}{(1-q\,\mathrm{e}^{-(1-q)T})^2} \ , \tag{3}$$

These results can be used directly to determine the statistics of branch lengths and the number of generations (birth events which have not led to an extant clade) along a branch of the phylogenetic tree. Of particular interest for the inference scheme presented in Section 2 is the joint probability of a branch and the number of generations along it, which we derive in the following.

#### B Joint probability of size-one clades and the number of generations

We measure time relative to the time of observation $T$ and define $\tau = T - t$. It is zero when the population is observed and increases as we look further into the evolutionary past.

We now derive the joint probability $p_1(i|\tau_s, \tau_e)$ for the following accumulation of events: a clade that started with a single individual at time $\tau_s$ has not died out by time $\tau_e < \tau_s$, none of the birth events in this time interval lead to additional extant clades at the time of observation $T$, and there are $i$ birth events along the resulting lineage between $\tau_s$ and $\tau_e$. Figure 1A illustrates the situation. The subscript 1 indicates that at time $\tau_e$ there is then only a single individual that may or may not have surviving offspring (or survive itself) at the time of observation. This extends the approach in (Dieselhorst and Berg 2024), where we derived the same probability but without considering the number of birth events. The birth events along a branch are the generations along that branch. Using the probabilities (2)-(3), $p_1(i|\tau_s, \tau_e)$ obeys the master equation

$$-\partial_{\tau_e} p_1(i|\tau_s, \tau_e) = -q p_1(i|\tau_s, \tau_e) - p_1(i|\tau_s, \tau_e) + 2p_0(\tau_e) p_1(i-1|\tau_s, \tau_e) \ . \tag{4}$$

The first term accounts for the death of the individual, the second term for the probability of a birth event (either leading to an additional observed lineage or to a $i+1$ hidden birth event). The last term adds the probability that the $i$-th birth event occurs and becomes 'hidden' in the phylogenetic tree, as one of the two descending clades dies out before observation. Defining the generating function $G(\mu, \tau_s, \tau_e) = \sum_{i=0}^{\infty} \mu^i p_1(i|\tau_s, \tau_e)$ yields

$$-\partial_{\tau_e} G(\mu, \tau_s, \tau_e) = -\left(q + 1 - 2\mu p_0(\tau_e)\right) G(\mu, \tau_s, \tau_e) \ , \tag{5}$$

which is easily solved by the separation of variables method.

In the case of pendant branches, we integrate from $\tau_s$ to $\tau_e = 0$ and get

$$\begin{aligned} G(\mu, \tau_s, \tau_e = 0) &= G(\mu, \tau_s, \tau_e = \tau_s) \times \exp\left\{-(1+q)\tau_s + \mu\left[2q\tau_s + 2\log\left(\frac{1-q}{1-q\,\mathrm{e}^{-(1-q)\tau_s}}\right)\right]\right\} \\ &= \exp\left\{-(1+q)\tau_s + \mu\left[2q\tau_s + 2\log\left(\frac{1-q}{1-q\,\mathrm{e}^{-(1-q)\tau_s}}\right)\right]\right\} \ . \end{aligned} \tag{6}$$

In the second line, we have used that $p_1(i|\tau_s, \tau_e = \tau_s)$ is one if $i = 0$ and zero otherwise (since in a vanishing time interval, there is no time for birth or death events) and therefore $G(\mu, \tau_s, \tau_e = \tau_s) = 1$. The resulting probability of a clade of given age $\tau_s$ having a single member at the time of observation after $i$ generations is

$$p_1(i|\tau_s, \tau_e = 0) = \frac{1}{i!}\partial_\mu^i|_{\mu=0} G(\mu, \tau_s, \tau_e = 0)$$
$$= \mathrm{e}^{-(1+q)\tau_s} \frac{1}{i!} \left[ 2q\tau_s + 2\log\left( \frac{1-q}{1-q\,\mathrm{e}^{-(1-q)\tau_s}} \right) \right]^i . \tag{7}$$

Summing over the number of generations results in equation (3).

Integrating the equation of motion of the generating function (5) only from $\tau_s$ to some intermediate $\tau_e > 0$, we find the joint probability of an individual having $i$ birth events in this time interval and all descendants of these births dying out by the time of observation

$$p_1(i|\tau_s, \tau_e) = \mathrm{e}^{-(1+q)(\tau_s-\tau_e)} \frac{1}{i!} \left[ 2q(\tau_s - \tau_e) + 2\log\left( \frac{1-q\,\mathrm{e}^{-(1-q)\tau_e}}{1-q\,\mathrm{e}^{-(1-q)\tau_s}} \right) \right]^i . \tag{8}$$

It can easily be checked that this solves the master equation (4) and simplifies to equation (7) for the case of $\tau_e = 0$.

For a given branch starting at $\tau_s$ and ending at $\tau_e$, this result implies that the number of generations is Poisson distributed with mean

$$\langle i \rangle(\tau_s, \tau_e) = 2q(\tau_s - \tau_e) + 2\log\left( \frac{1-q\,\mathrm{e}^{-(1-q)\tau_e}}{1-q\,\mathrm{e}^{-(1-q)\tau_s}} \right) . \tag{9}$$

This agrees with the result derived by Bokma, van den Brink, and Stadler (Bokma et al. 2012) for the mean number of generations. The first term in (9) is proportional to the elapsed time $\tau = \tau_s - \tau_e$; the constant of proportionality is twice the relative rate of cell death. Without death events, every birth event would produce two extant lineages, thus producing an interior node in the tree. As a result, without cell death ($q = 0$), the number of generations along each branch is zero. The second, nonlinear term is zero at $\tau = 0$ and grows with increasing branch length. This means that the mean number of generations along a branch is not proportional to the length of the branch calendar time. This counterintuitive result has a simple explanation: all extra lineages arising by birth processes along a branch must die out by the time the population is sampled, and this happens with a smaller probability at short $\tau$, reducing the number of generations for short branches. The effect decreases with increasing $\tau_e$.

Figure 1B compares these results to numerical simulations. A population with relative death rate $q = 3/4$ is grown from a single individual to 7500 individuals (stopping the simulation when this population size is reached) and the phylogenetic tree is constructed. Figure 1B shows the mean and variance of the number of generations along pendant branches of a given length $\tau$. Crucially, the mean number of generations is clearly not proportional to the branch length $\tau$, but follows (9) for $\tau_e = 0$. (For a Poisson distribution, the variance equals the mean.)

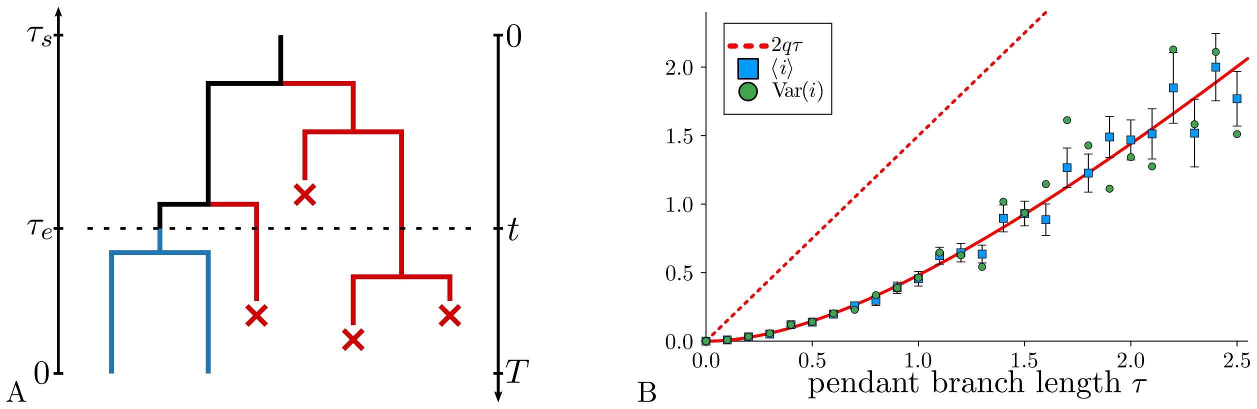


**Figure 1: Generations along a branch and the distribution of the number of generations along pendant branches.** (A) This schematic illustrates generations along a branch, which arise from birth events where one of the clades generated by the event dies out by the time of observation. Along the branch between $\tau_s$ and $\tau_e$ there are $i = 2$ such generations, with the clades going extinct shown in red. (B) Numerical results come from a single population grown to size 7500 at relative death rate $q = 3/4$. We pick all pendant branches of a given length $\tau$ ($\pm 0.05$) and plot the mean (large blue symbols, with the standard error given by the error bars) and variance (small green symbols) of the number of generations along these branches against $\tau$. The full red line gives the theoretical result (9) (with $\tau_e = 0$) for both the mean and the variance. The dotted red line gives the linear term $2q\tau$ in (9), corresponding to generations appearing along the branch at a constant rate.

Summing over the number of generations $i$ in distribution (8) gives the distribution of branch lengths $p_1^{(\tau_e)}(\tau_s - \tau_e)$ (Dieselhorst and Berg 2024). Equation 8 can also be derived by multiplying $p_1^{(\tau_e)}(\tau_s - \tau_e)$ with the result from (Bokma et al. 2012) that the number of generations is Poisson distributed with mean (9).

These results can be easily adapted to finite sampling as discussed in Section 2.2. $p_0(\tau_e)$ in the equation of motion (5) for the generating function needs to be replaced by $p_0^{(\rho)}(\tau_e)$ given by (6). The result is (8) but with the term in square brackets replaced by

$$\langle i \rangle^{(\rho)}(\tau_s, \tau_e) = 2q(\tau_s - \tau_e) + 2 \log \frac{(1 - \rho - q)\, \mathrm{e}^{-(1-q)\tau_e} + \rho}{(1 - \rho - q)\, \mathrm{e}^{-(1-q)\tau_s} + \rho} \,, \tag{10}$$

which is the mean number of generations on a branch running from $\tau_s$ to $\tau_e$ generalized to finite sampling. In the iterative algorithm of Section 2.1, each leaf of the tree is assigned an additional factor of $\rho$ accounting for sampling at that probability.

### C Distribution of the number of mutations along a branch

Under a constant molecular clock hypothesis, the number of mutations along a branch of a given length is Poisson distributed with a mean given by the branch length (in calendar time) multiplied by a mutation rate per time.

However if, under cellular reproduction, mutations occur at cell divisions, different statistics of the number of mutations along a branch arise. As we have seen in the previous section, the number of generations along a given branch follows a Poisson distribution. If the number of mutations per cell division is itself a random variable this makes the total number of mutations along a branch follow a compound Poisson distribution: The number of accumulated mutations $m$ is the sum over $i$ independent, identically distributed random variables $\hat{m}$, where the number of generations $i$ itself is a Poisson distributed random variable.

Denoting with $i$ the random variable of the number of generations along a branch from $\tau_s$ to $\tau_e$ and with $\hat{m}$ the (also random) number of mutations at a birth event, we can use Wald's equation (Ross 2014) to compute the mean number of mutations between bifurcations as

$$\langle m \rangle = \langle i \rangle \langle \hat{m} \rangle \ . \tag{11}$$

For simplicity, we here suppress the dependencies of $i$ and $m$ on $\tau_s$ and $\tau_e$. Using the law of total variance (also known as Eve's law, (Blitzstein and Hwang 2019)), we find the variance to be

$$\begin{aligned} \mathrm{Var}\,[m] &= \langle \mathrm{Var}\,[m|i] \rangle + \mathrm{Var}\,[\langle m|i \rangle] \\ &= \mathrm{Var}\,[\hat{m}]\,\langle i \rangle + \langle \hat{m} \rangle^2 \mathrm{Var}\,[i] \\ &= \left( \mathrm{Var}\,[\hat{m}] + \langle \hat{m} \rangle^2 \right) \langle i \rangle \ . \end{aligned} \tag{12}$$

When we take the number of mutations per birth event to be Poisson distributed with $\langle \hat{m} \rangle = \mathrm{Var}\,[\hat{m}] = \mu$, we obtain

$$\mathrm{Var}\,[m] = \langle i \rangle \mu (1 + \mu) \tag{13}$$

which differs by a factor of $(1+\mu)$ from the corresponding result for Poisson-distributed accumulated mutations. The probability mass function for the mutations along a given branch is

$$P(m|\tau_s, \tau_e) = \sum_{i=0}^{\infty} P(i|\tau_s, \tau_e) P(m|i) \tag{14}$$

$$= \sum_{i=0}^{\infty} \mathrm{e}^{-\langle i \rangle (\tau_s, \tau_e)} \, \frac{\langle i \rangle (\tau_s, \tau_e)^i}{i!} \, \mathrm{e}^{-i\mu} \, \frac{(i\mu)^m}{m!} \ . \tag{15}$$

$P(i|\tau_s, \tau_e)$ denotes the Poisson distribution for the number of generations along a given branch from $\tau_s$ to $\tau_e$.

Note that here we focus on the mutations accumulating *between* two tree bifurcations and have not counted the mutations arising at the cell division establishing the branch. For this reason, $P(m|i)$ is a regular Poisson distribution, whereas (2) features an additional generation at the top of the branch. (For this reason, in the computation of the likelihood as described in Section 2, we used the compound Poisson distributed mutations along the branch plus a Poisson distributed number of mutations from the birth event starting the branch.)

Two examples of the probability mass function (14) are shown in Figure 2. On the left, we show the distribution for $\langle i \rangle = 10$ and $\mu = 1$ and on the right for $\langle i \rangle = 3$ and $\mu = 10$ (red lines). The Poisson distributions with the same mean $\langle i \rangle \mu$ are depicted by grey dashed lines.

#### D Comparison of models with different mutation statistics

By using the statistics of the number of generations along branches, our approach incorporates two differences to strict molecular clock models: The mean number of generations along branches is inhomogeneous throughout the tree (as discussed in Supplementary Information B), which affects the statistics of mutations if they are tied to birth events. Furthermore, mutations from birth events along branches follow compound Poisson statistics rather than a Poisson distribution (Supplementary Information C).

In order to compare approaches based on different assumptions on the mutational process, we have implemented the same likelihood-based inference scheme introduced above but with the assumption that mutations occur at some rate per time. This yields a Poisson distribution for the number of mutations along a given branch. We do this in two different ways. In a first approach, we take the number of mutations along a branch to be Poisson distributed, with a mean depending on the number of generations. Specifically, the mean number of mutations along a branch running from $\tau_s$ and $\tau_e$ is taken to be $(1 + \langle i \rangle(\tau_s, \tau_e)) \times \mu$, i.e. the expectation value of the number birth events contributing mutations to the branch (one birth event forming the branch plus the mean number of generations along the branch $\langle i \rangle(\tau_s, \tau_e)$) times the mean number of mutations per birth event $\mu$. This replaces the compound Poisson statistics (see Supplementary Information C) by a Poisson distribution with the same mean. In a second approach, we assume a constant molecular clock with rate $\mu_\mathcal{T}$ throughout the entire tree. In both approaches, the bifurcation times and number of generations along branches are integrated and summed over as discussed in Section 2.1.

To compare the inference results of these different models, we simulated trees with relative death rate $q = 3/4$ and mean mutations per birth event $\mu = 1$. In Figure 2A, we show the inferred parameters from 100 populations, where we stopped the simulations when a population size of $N = 300$ was reached. For Figure 2B (and C), we stopped when the population reached size $3 \times 10^3$ $(3 \times 10^4)$ and reconstructed the tree from 300 randomly chosen individuals (corresponding to sampling probability $\rho = 0.1$ or $\rho = 0.01$, respectively). Violin plots show the distributions of the inferred values of $q$ for the different models of the number of mutations along each branch. On the left of each subfigure, we show the inference results for the original likelihood (5) based on the compound Poisson statistics, in the centre we use the Poisson statistics with the same mean, and on the right the constant-rate molecular clock. The three histograms below depict the corresponding distributions of the inferred mutation parameters $\mu$ and $\mu_\mathcal{T}$.

In the case of complete sampling, we find that the first approach (Poisson statistics) gives results

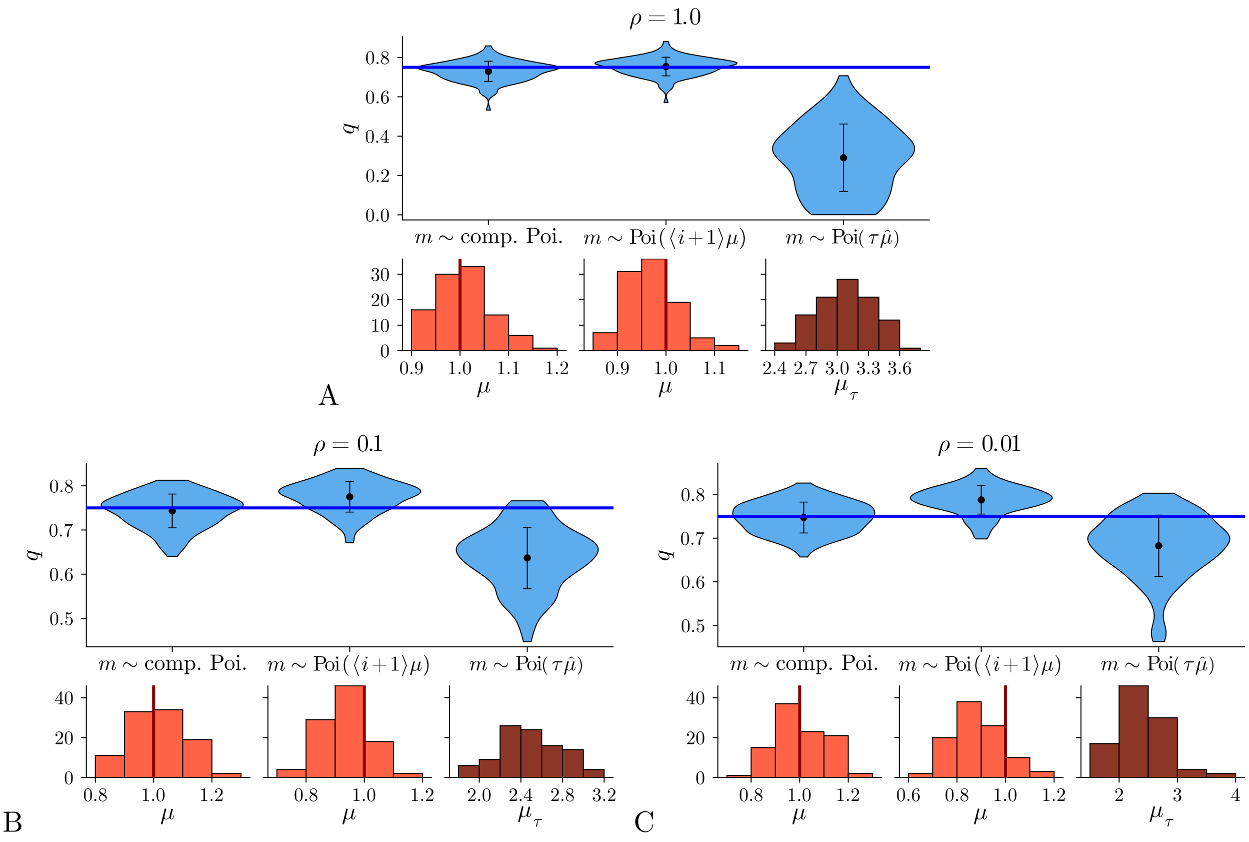


**Figure 2: Comparison of different models of mutations.** We compare results of our method based on the compound Poisson statistics of mutations along branches ("$m \sim$ comp. Poi.") and the same algorithm but with Poisson statistics (inhomogeneous "$m \sim \mathrm{Poi}(\langle i+1 \rangle \mu)$" and a constant-rate molecular clock "$m \sim \mathrm{Poi}(\tau\mu_\tau)$", see text). (A) Distributions of the inferred parameters from 100 populations grown to size 300. Violin plots show the results for $q$, with the dots indicating the mean and the error bars the standard deviation. Histograms show the inferred mean number of mutations per cell division $\mu$ or mutation rate $\mu_\tau$. Solid lines show the underlying parameter values $q = 3/4$ and $\mu = 1$. (B) The same, but for sampling probability $\rho = 0.1$. (C) The same for $\rho = 0.01$.

comparable to the compound Poisson statistics. However, in the cases of $\rho = 0.1$ and $\rho = 0.01$, a noticeable bias is seen in the results based on Poisson statistics. This is because in the fully sampled case, many (in particular pendant) branches have no hidden generations along the branch. The mutations solely come from the birth event establishing the branch and the number of mutations is thus Poisson distributed. When only a fraction of the extant distribution is sampled, the number of hidden birth events (in particular along pendant) branches increases, and thus, the mutations follow the compound Poisson rather than Poisson statistics.

While the inferred values of $q$ using the molecular clock model are inaccurate in all three cases, a small improvement is seen when comparing the results under finite sampling to the ones from fully observed populations (note the different $y$-axes). This is because the dependence of the mean number of generations on the age of a branch (temporal inhomogeneity) becomes weaker for older branches. Generally, the mean number of generation per branch given by equation (1) depends both on $\tau_s$ and $\tau_e$. For large $\tau_s$ and $\tau_e$, however, it asymptotically depends only on the difference $\tau_s - \tau_e$, as it would under a molecular clock. For this reason, with decreasing sampling probability, the effects of the temporal inhomogeneity decrease as more branches lie further away from the time of observation. This produces a small improvement in the inference results under the molecular clock assumption for $\rho = 0.01$ compared to $\rho = 0.1$.

#### E Inference of the sampling probability

In Section 2.2, we discussed the inference of the sampling probability $\rho$ together with the other parameters $q$ and $\mu$. We illustrate this inference by simulating populations with $q = 3/4$ and $\mu = 1$, stopping when the population size reaches 3000. We then reconstruct the phylogenetic trees based on 300 randomly chosen extant individuals, which corresponds to a sampling probability of $\rho = 0.1$. We created 100 trees in this manner and infer the corresponding parameters. Figure 3A shows the inference results for $q$ and $\mu$ at given $\rho = 0.1$, while Figure 3B depicts the results of the simultaneous inference of all three parameters. As expected, $q$ and $\mu$ are more accurately inferred (with a lower standard deviations (std) in particular in the inferred $\mu$) when $\rho$ is known. However, it is still possible to estimate all three parameters.

#### F Inference bias

In Figure 3 of the main text we showed how model parameters can be inferred from simulated data for a specific choice of model parameters. Here we repeat the inference from artificial data trees of size $N = 300$ across a wide range of model parameters $q$ and $\mu$ and for both full ($\rho = 1$) and finite ($\rho = 0.01$) sampling. The resulting inference error of the model parameters remains small and nearly flat over the entire range of values for $q$, see Figure 4A and C. In Figure 4B and D, we show that the accuracy of the maximum likelihood estimates improves significantly for higher $\mu$. As $\mu$ decreases the inference becomes less reliable as then only few generations are actually marked by

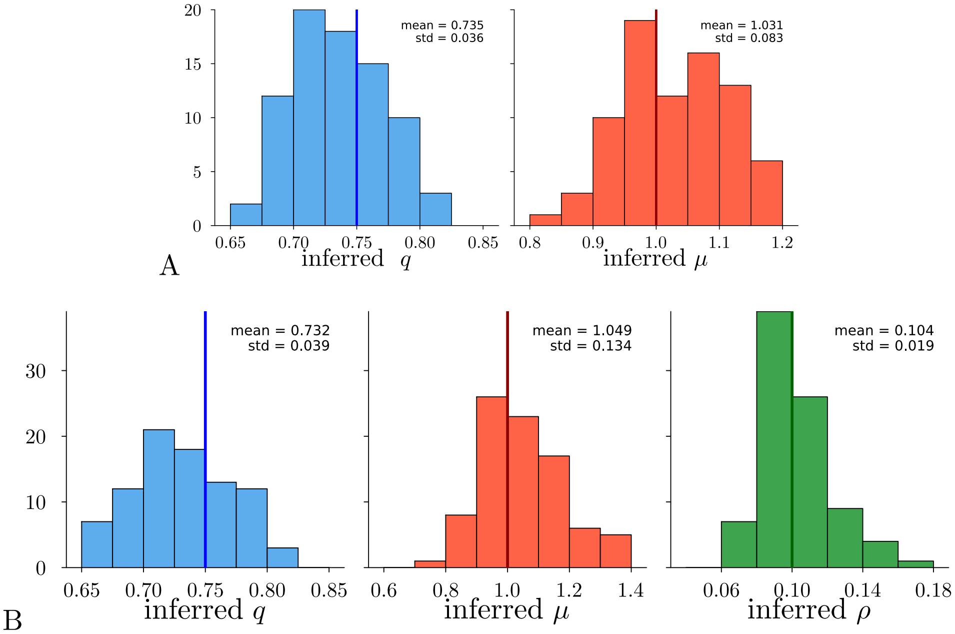


**Figure 3: Parameter inference from samples of a population.** Depicted are the inference results from 100 populations simulated with $q = 3/4$ and $\mu = 1.0$ up to size 3000. For each population, a tree was reconstructed from 300 samples, corresponding to a sampling probability of $\rho = 0.1$ (see text). (A) Inference results when the value of $\rho = 0.1$ is known. (B) Inference results when $\rho$ is inferred along with the other parameters.

mutations. In this regime, the bottleneck of the analysis will be the construction of the phylogeny, which is outside the our scope here.

Maximum likelihood estimates generally have a bias in the parameter estimates (Cover and Thomas 2006), which vanishes in the limit of a large data. Figure 5 shows how the bias in the parameter estimates from the likelihood (5) decreases with the number of sampled individuals (equal to the population size at full sampling, $\rho = 1$). This bias can be reduced by conditioning the population e.g. on not dying out. Such conditions are applied by replacing the uniform prior in the integration over the time of the root node (5). We refer to (MacPherson et al. 2022) for a discussion on different conditioning scenarios.

#### G Combining different mutational processes

So far we have focussed on mutations that are caused by the reproduction of cells and arise at each cell division with a certain probability. Also in somatic evolution mutations can arise per unit time, or more precisely, DNA lesions are generated at a certain rate per unit time, and lead to mutations unless they have been repaired by the time the cell divides (Spisak et al. 2023). In this section, we extend our model of mutations to the case where mutations occur both at a constant rate $\mu_\mathcal{T}$ per unit time (i.e. following a molecular clock) and, in addition, each division contributes a number of mutations drawn from a Poisson distribution with mean $\mu$ as described in Section 2.1.

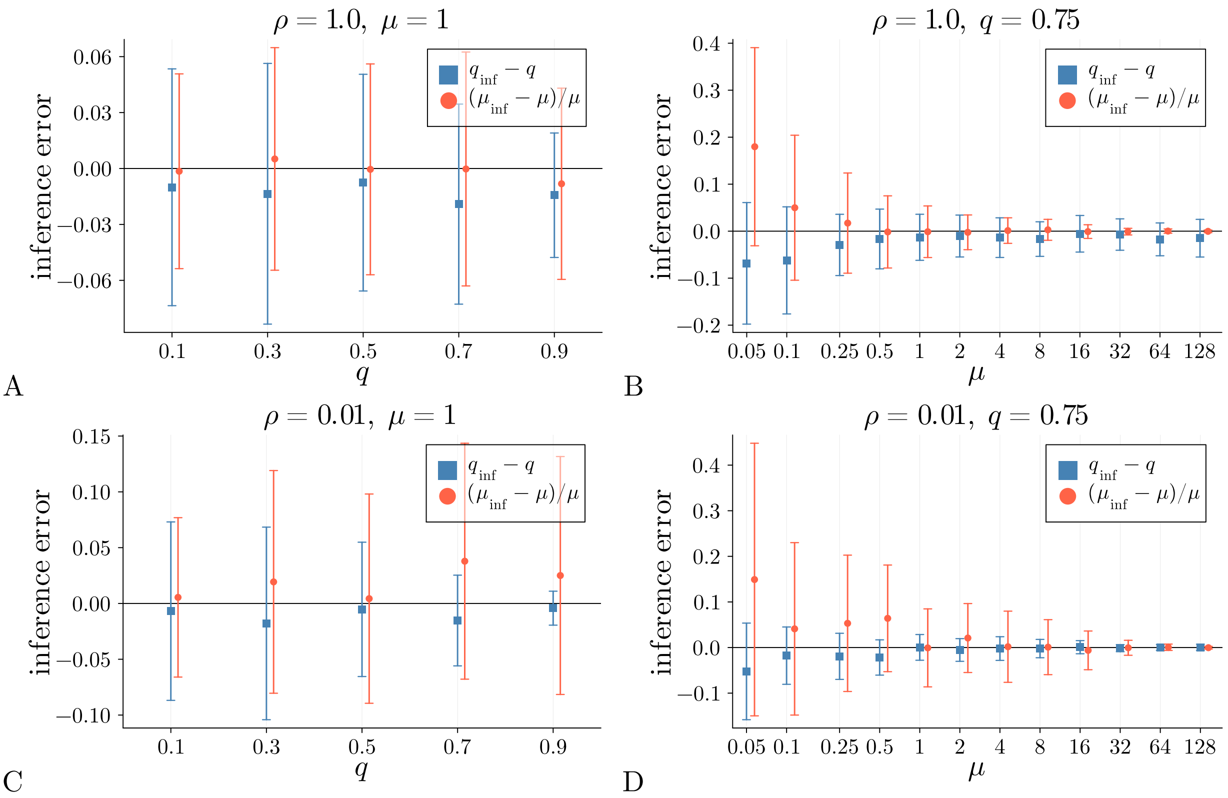


**Figure 4: Inference error across different values of the model parameters.** The difference between inferred and underlying values of the relative rate of death $q$ and the number of mutations per birth $\mu$ is shown for different values of the underlying $q$ and $\mu$. In case of $\mu$, the difference relative to the true values is shown. The symbols (blue squares for $q$ and orange circles for $\mu$) show the differences between inferred and underlying values averaged over 100 runs each producing a tree of size $N = 300$. Error bars show the corresponding standard deviation. The inference is based on maximizing the likelihood (5). For visibility, the symbols indicating the results for the mutation parameter $\mu$ are shifted a little to the right. (A) Results from fully sampled populations with the mean number of mutations kept fixed at $\mu = 1$, while the relative rate of death $q$ varies as shown on the $x$-axis. (B) The same for varying $\mu$ (as shown on the $x$-axis) and relative rate of death fixed at $q = 0.75$. (C)-(D) The same as (A)-(B), but for a finite sampling probability of $\rho = 0.01$. In each simulation, a population was grown to size 30000 and trees were built from 300 randomly chosen individuals.

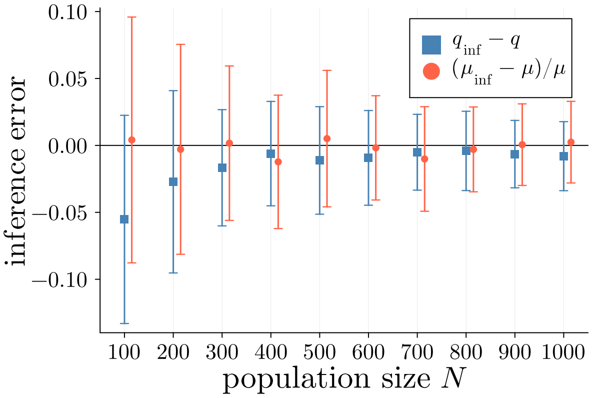


**Figure 5: The dependence of the inference error on population size.** The difference between inferred and underlying values of the relative rate of death $q$ and the number of mutations per birth $\mu$ is shown for populations of different sizes. In case of $\mu$, the difference relative to the true values is shown. The symbols (blue squares for $q$ and orange circles for $\mu$) show the averages over 100 runs; error bars show the corresponding standard deviation. The underlying parameters were $q = 3/4$, $\mu = 1$ at full sampling ($\rho = 1$). As expected, the bias of the estimates decreases with the population size $N$, until reaching a plateau for populations of size $N = 400$ and larger.

Mutations that occur at a fixed rate $\mu_\tau$ per unit time lead to a Poisson distributed number of mutations with mean $\mu\tau$ along a branch of length $\tau$. The total number of mutations arising both over time and from $i$ birth events is thus

$$P(m|i,\tau) = \exp(-\mu_\tau\tau - (i+1)\mu)(\mu_\tau\tau + (i+1)\mu)^m/m! \; . \tag{16}$$

Using this distribution in equations (3)-(5), we can compute the likelihood of $\mu_\tau$ along with $q$ and $\mu$.

However, teasing apart the relative contributions of these two processes from a phylogenetic tree will require larger trees with more mutations along the branches than currently available and must be left for future work. Relying on the mutational signatures (Spisak et al. 2023) to tell apart $m_\tau$ (the number of mutations generated a process running per unit time) and $m - m_\tau$ (the number of mutations generated by a process arising at cell divisions) would simplify the inference. We note that if the number of mutations in a birth event is not Poisson distributed, an additional sum arises when computing the distribution of mutations (16).

Finally, we look at sequencing noise, modelled by adding a random number $m_e$ of sequencing errors (amplification errors) to the tree leaves. These numbers are drawn independently and identically from some distribution $P_e(m_e)$. Incorporating this model into the iterative construction of the likelihood leads in place of (3) to

$$P(\nu|\tau_s) = \sum_{m_e=0}^{m} P_e(m_e) \sum_{i=0}^{\infty} P(m - m_e|i) p_1(i|\tau_s, \tau_e = 0) \; . \tag{17}$$

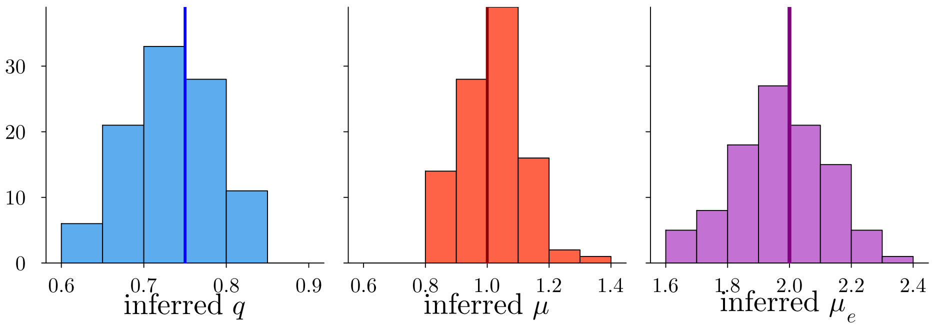


**Figure 6: Inference with sequencing error model** (17) **describing erroneous additional mutations.** We show the inference results from 100 fully sampled populations of size 300, simulated with $q = 0.75$ and $\mu = 1$. In order to mimic sequencing errors, pendant branches were extended by a Poisson distributed number of mutations with mean $\mu_e = 2$, see text. The distributions of inferred values for all three parameters are depicted by the histograms. Vertical lines indicate the true underlying parameter values.

We take the natural choice that the distribution $P_e(m_e)$ is a Poisson distribution with mean $\mu_e$ to be inferred. In this case, similar to the case with mutations accumulating over time, performing the sum over $m_e$ explicitly can be avoided. However, other distributions can be implemented as well. The statistics of interior branches is not affected by erroneous additional mutations. Hence, the iterative procedure (4) for the interior branches remains unchanged, and the model parameters including $\mu_e$ can be inferred by maximizing the resulting likelihood. We show the inference results from 100 populations simulated under this model in Figure 6.

Another source of sequencing errors — in particular in single-cell sequencing — is so-called allelic dropout, where individual nucleotides fail to be sequenced for technical reasons. This means loci that are really heterozygous appear homozygous, and the mutation remains undetected. The dropout probability depends on the specific sequencing technology, but recent technologies consistently yield dropout probabilities per nucleotide below 25% (Estévez-Gómez et al. 2025).

Mutations which remain undetected in multiple samples will affect the reconstruction of the phylogenetic tree and the assignment of mutations to specific branches. In order to quantify the inference bias arising from allelic dropout under the infinite sites assumption, we have simulated phylogenetic trees and removed each mutation in each extant individual independently with probability $\sigma$. Mutations were then assigned to the branch above the most recent common ancestor of all individuals carrying the mutation (or the pendant branch above a single individual, if a mutation is found nowhere else). Thus, with increasing dropout probability $\sigma$ mutations are moved to successively lower branches of the tree or removed entirely. We then inferred the model parameters $q$ and $\mu$ from altered assignment of mutations to branches. Figure 7 A-D shows the inference results for different values of $\sigma$. We find that for dropout probabilities $\sigma$ below 0.25 the value of $q$ is underestimated while the sign of the bias of $\mu$ depends on its true underlying value and the dropout probability $\sigma$. However, for the range of dropout probability investigated here, the relative errors remained below 15%. In Figure 7 E-H, we repeated the inference from trees with $\sigma = 0.25$ but under the assumption of different mutation statistics as introduced in Supplementary Information D. As seen before without the allelic dropout, we find that the inferred values under a strict molecular clock assumption (violin plots on the right of the figures) vary widely and are inaccurate. Interestingly, the approach with Poisson statistics for the number of mutations along branches with a mean proportional to the mean number of generations (violin plots in the center) seems relatively unaffected by allelic dropout, yielding results comparable to the ones in Supplementary Information D: the inference results are reliable under full sampling, but show a bias under finite sampling.

As discussed in Section 3, future extensions of our approach can deal with uncertainty in the reconstructed tree by summing over multiple trees compatible with the sequences of extant individuals. In this manner, allelic dropout could be explicitly accounted for in the inference scheme. However, with further advancements in single-cell sequencing, allelic dropout is expected to become less of an issue in the future. We also note that the stem cell data from (Mitchell et al. 2022) used in Section 2.3 is based on sequencing small colonies derived from single-cells. This yields much lower levels sequencing noise compared to sequencing single cells, and almost negligible dropout probabilities and amplification errors (Miao et al. 2020).

#### H Inference of selection

The algorithm presented in Section 2.1 can easily be adapted to account for distinct parameters in different clades of the tree. This enables detecting mutations which change the evolutionary dynamics in the corresponding clade. In principle, each such parameter change leads to three additional parameters to infer: The birth and death rates of the distinct subtree, $\beta_{\mathrm{sub}}$ and $\delta_{\mathrm{sub}}$ respectively, in units of the main tree's birth rate and the mean number of mutations per generation $\mu_{\mathrm{sub}}$ in the subtree. In the following, we will consider the case where there is only a single such parameter change, leading to a single distinct subtree.

Taking the parameter change to occur at a particular node in the tree (with both descending lineages following the new parameters), we can easily compute the probability of the distinct subtree using the iterative algorithm from Section 2.1, replacing $\mu$ in equation (2) with $\mu_{\mathrm{sub}}$ and using

$$p_1(i|\tau_s,\tau_e) = \mathrm{e}^{-(\beta_{\mathrm{sub}}-\delta_{\mathrm{sub}})(\tau_s-\tau_e)}\,\frac{1}{i!}\left[2\delta_{\mathrm{sub}}(\tau_s-\tau_e) - \log\frac{\beta_{\mathrm{sub}}-\delta_{\mathrm{sub}}\,\mathrm{e}^{-(\beta_{\mathrm{sub}}-\delta_{\mathrm{sub}})\tau_e}}{\beta_{\mathrm{sub}}-\delta_{\mathrm{sub}}\,\mathrm{e}^{-(\beta_{\mathrm{sub}}-\delta_{\mathrm{sub}})\tau_s}}\right]^i \tag{18}$$

in equations (3) and (4). All iterations over clades $\nu$ starting with branches which are not part of the distinct subtree are performed as before with parameters $q$ and $\mu$. The computation of $P(\nu|\tau_s)$ (equation (4)) for the branch above the parameter change also uses these main parameters but uses the results of the two descending clades $P(\nu'|\tau_s)$ and $P(\nu''|\tau_s)$ which were computed with the alternative parameters. Proceeding until the final integration over the root node (5), we obtain the probability of the entire tree with a distinct subtree, and hence the likelihood of parameters $q$, $\mu$, $\beta_{\mathrm{sub}}$, $\delta_{\mathrm{sub}}$ and $\mu_{\mathrm{sub}}$.

Proposing parameter changes at different nodes of the tree and comparing the resulting maximum

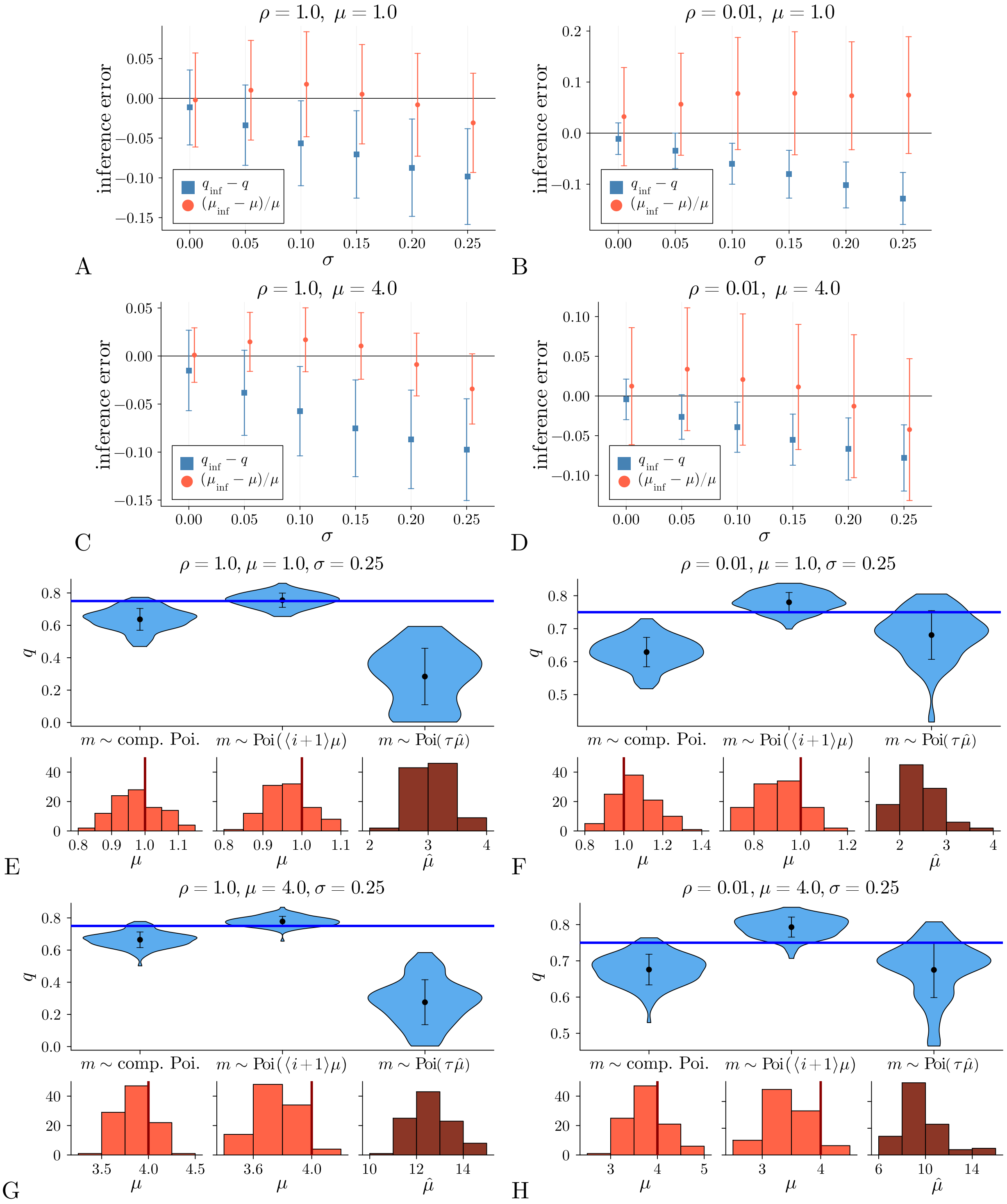


**Figure 7: Inference bias arising from allelic dropout.** A-D: We simulated 100 phylogenetic trees with 300 leaves each and a relative rate of death $q = 0.75$. The mean number of mutations per generation $\mu$ and the sampling probability $\rho$ are indicated in the figure titles. We then simulated allelic dropout as described in the text with different dropout probabilities $\sigma$ shown along the $x$-axis. We then inferred $q$ and $\mu$ from the altered assignment of mutations to branches (see text) and computed the deviation of $q$ and relative deviation of $\mu$ from the true underlying parameters. The symbols (blue squares for $q$ and orange circles for $\mu$) show the inference error averaged over the 100 trees; error bars show the corresponding standard deviation. E-H: We compare the inference results of trees with allelic dropout of $\sigma = 0.25$ under a different mutation statistics where mutations arise along branches at a fixed rate per unit time, see Supplementary Information D.

likelihoods allows finding the most likely location of a parameter change. To test this approach, we have simulated 100 trees in the following way: We simulate a population starting with a single individual and with parameters $q = 0.75$ and $\mu = 1$. The two lineages descending from the $200^{\text{th}}$ birth event evolve under a higher birth rate $\beta_{\text{sub}} = 2$, but with otherwise identical parameters $\delta_{\text{sub}} = 0.75$ and $\mu_{\text{sub}} = 1$. The simulation is stopped when the population first reaches a size of $N = 300$ and the phylogenetic tree is reconstructed from all extant individuals. If the population has died out or either of the two distinct parts of the tree have less than 50 leaves, the simulation is repeated.

We then inferred parameters under the assumption of a single parameter change. This parameter change was proposed at every node where the descending subtree includes at least 25 leaves. For each of these proposed parameter changes, the likelihood was determined. We fixed the discretization of time to 1000 equidistant subintervals between 0 and $2\log(N/\rho)/(1 - q_{\text{homo}})$, where $q_{\text{homo}}$ is the maximum likelihood estimate of the relative death rate under a homogeneous parameter model. The upper limit of the sum over the number of generations was set to 50. Comparing the results of each proposed parameter change, we accepted the one with the highest likelihood maximum.

In order to assess the accuracy with which we have found the correct location of the parameter change, we define the dichotomy error as follows: If the inferred distinct subtree of size $n_{\text{inf}}$ overlaps with the true distinct subtree of size $n_{\text{true}}$, the dichotomy error is given by $n_{\text{inf}} - n_{\text{true}}$. If the two do not overlap, the dichotomy error is given by the number of nodes which is in neither of the two subtrees. In our simulations of trees with 300 leaves (and therefore 599 nodes), this dichotomy error is given by $599 - n_{\text{inf}} - n_{\text{true}}$.

Figure 8 shows the results from 100 simulated populations. In 58 cases, the parameter change was detected at the correct position. Only in four runs did the inferred subtree not overlap with the true distinct subtree. The dichotomy errors are depicted in Figure 8A, the inferred parameters from all 100 simulations are plotted as violin plots in Figure 8B. We also show the inferred relative death rate $q_{\text{sub}} = \delta_{\text{sub}}/\beta_{\text{sub}}$. Figure 8C shows that the results of only the simulations where the subtree was found correctly are only slightly better than when all runs are included. The selective advantage of the distinct subtree, i.e. a smaller relative death rate $q_{\text{sub}} < q$, was inferred correctly in all 96 cases where the inferred subtree overlapped with the true distinct subtree.

So far, we have simply used the log-likelihood difference as a rule to accept or reject different proposed parameter changes. However, we find that propositions of small distinct subtrees can be accepted due to overfitting stochastic fluctuations. This is why, here, we only consider subtrees with 25 leaves or more. Future research could lead to a principled rule for accepting/rejecting parameter changes that takes into account such fluctuations.

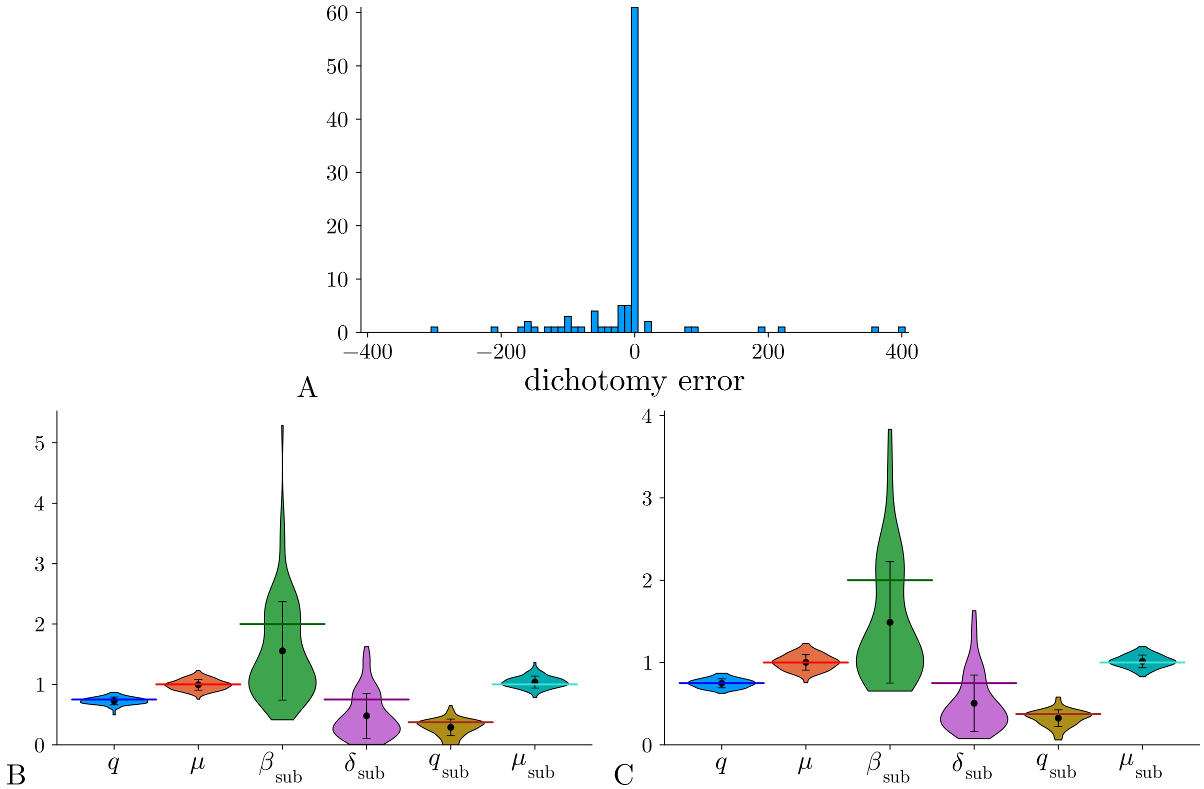


**Figure 8: Results of the inference of parameter shifts** We show inference results from 100 simulated populations as described in the text. (A) Dichotomy errors of the inferred distinct subtrees, as defined in the text. The correct subtree (dichotomy error zero) was found in 58 of the 100 simulations. (B) Violin plots show the inferred parameters, with black dots marking the mean and error bars the standard deviations of the inference results. Horizontal lines mark the true values used in the simulation. We further depict the relative death rate in the subtree $q_{\text{sub}} = \delta_{\text{sub}}/\beta_{\text{sub}}$. (C) The same as (B), but only depicting the results of the 58 simulations where the distinct subtree was located correctly.

### I Inference from prenatal HSC data under molecular clock assumption

In Section 2.3.1, we inferred the relative death rate $q$ and the mean number of mutations $\mu$ for trees of haematopoietic stem cells (HSC) sampled from cord blood of neonates (donor CB001 and CB002 of (Mitchell et al. 2022)). Here, we give details on the inference under the assumption of a molecular clock with mutation rate $\mu_\mathcal{T}$. This is done for comparison with the model (2), where mutations occur at cell divisions. In Figure 9, we show the log-likelihood landscape of $q$ and $\mu_\mathcal{T}$ for donor CB001 and CB002. The maximum of the likelihood is found at $q = 0$ and $\mu = 6.807$ for donor CB001 and at $q = 0$ and $\mu = 6.89$ for donor CB002. In Section 2.3.1, we argued that a vanishing relative death rate is implausible because HSC differentiate to other cell types and are thus removed from the population of HSC, which should result in a finite value of $q$. Here, we derive the confidence regions around the maximum likelihood point estimates in order to better evaluate the results under the molecular clock assumption.

Standard approaches to estimate a confidence interval cannot be applied when the likelihood maximum is at the boundary of parameter space. To estimate a confidence region in this case, we assume uniform prior probabilities distributions for $q \sim \mathcal{U}(0, 1)$ and $\mu_\mathcal{T} \sim \mathcal{U}(0, 10)$. In this case, the likelihood depicted in Figure 9 is proportional to the joint posterior probability density of $q$ and $\mu_\mathcal{T}$. We can then find Bayesian credibility regions by computing the highest posterior density (HPD) regions for a given percentage of the probability mass. The results are indicated by red contour lines in Figure 9. We find that for donor CB001 the 95% HPD region includes values of $q$ up to 0.13 for $\rho = 0.01$ and up to 0.26 for $\rho = 0.001$. For donor CB002, the upper bound is found at $q = 0.08$ when setting $\rho = 0.01$ and at $q = 0.16$ for $\rho = 0.001$. Since this analysis uses a uniform prior rather than a fully specified Bayesian model, it provides only an approximate characterization of uncertainty under the molecular clock assumption.

To estimate the relative rate of differentiation (or cell death) of HSCs during development independently of a model of mutations, we use *in vivo* measurements of HSC population size. Ema and Nakauchi (Ema and Nakauchi 2000) reported that the absolute number of HSC increases approximately 38-fold between embryonic day 12 (E12) and E16. Assuming an average cell-cycle time of $12\,h$, this corresponds to roughly 8 HSC divisions during this window, and a relative rate of differentiation/death of $q = 0.35$. This is only a rough estimate, which is likely influenced by subpopulations of HSC differentiating at different rates, among other factors.

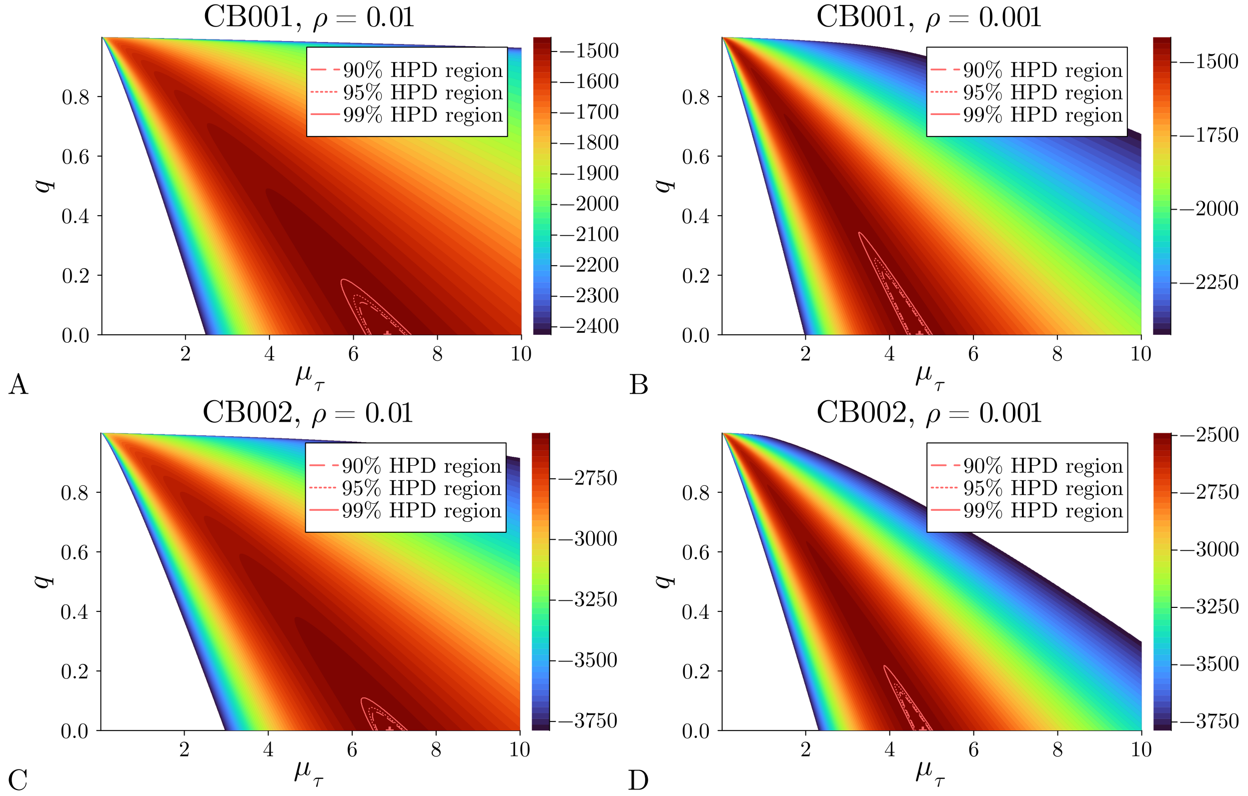


**Figure 9: Parameter likelihood under molecular clock assumption.** We show the log-likelihood landscape of the relative death rate $q$ and mutation rate $\mu_\tau$ for the trees of HSC from donors CB001 and CB002 from (Mitchell et al. 2022) and for sampling probabilities $\rho = 0.01$ and $\rho = 0.001$. The likelihood maxima are indicated by a red cross (only half is shown). The red contours indicate parameter regions enclosing specified posterior percentiles, assuming the likelihood is proportional to the posterior (see text).
